# Quantifying male and female pheromone-based mate choice in *Caenorhabditis* nematodes using a novel microfluidic technique

**DOI:** 10.1101/196733

**Authors:** Flora Borne, Katja R. Kasimatis, Patrick C. Phillips

**Affiliations:** Ecole Normale Supérieure Paris-Saclay, Université Paris-Saclay, Cachan, France; Institute of Ecology and Evolution, University of Oregon, Eugene, Oregon, United States of America

**Author notes:** Corresponding author (PCP). These authors contributed equally to this work (Flora Borne, Katja R. Kasimatis).

## Abstract

Pheromone cues are an important component of intersexual communication, particularly in regards to mate choice. *Caenorhabditis* nematodes predominant rely on pheromone production for mate finding and mate choice. Here we describe a new microfluidic paradigm for studying mate choice in nematodes. Specifically, the Pheromone Arena allows for a constant flow of small molecule signals to be passed in real time from signaling worms to those making a choice without any physical contact. We validated this microfluidic paradigm by corroborating previous studies in showing that virgin *C. remanei* and *C. elegans* males have a strong preference for virgin females over mated ones. Moreover, our results suggest that the strength of attraction is an additive effect of male receptivity and female signal production. We go on to explicitly examine female choice and find that females are more attracted to virgin males. However, a female’s mate choice is strongly dependent on her mating status.

## Introduction

A critical component of sexual reproduction is the ability to find and recognize the appropriate individual with which to mate. Individuals must be able to distinguish members of their own species – namely, conspecifics – from those of other species – heterospecifics. Perhaps equally important, is the ability to choose high quality individuals that are receptive to mating. The process of mate choice is shaped by sexual selection and relies on communication between the sexes [1-3]. In particular, sex pheromones – small chemicals produced by a signaler to induce a sexual response in a receiver – are a major means of intersex communication across a wide variety of both invertebrate and vertebrate taxa [reviewed in 4]. Some of the best studied sex pheromones are the cuticular hydrocarbon family found in many insect species [5,6]. These pheromones have both species-specific and sex-specific effects [6]. For example, in *Drosophilae* female hydrocarbons attract males, while male hydrocarbons have an anti-aphrodisiac effect on other males and increase female receptivity [7-9]. While these studies highlight the importance of pheromones in mate choice, many of the taxa studied have additional behaviors and traits that contribute to the mate choice process, potentially confounding the relative reliance on pheromone-based cues.

*Caenorhabditis* nematodes rely almost exclusively on pheromone signals for mate choice [10,11]. Pheromone signals in *Caenorhabditis* have traditionally been studied using plate-based chemotaxis assays, where male attraction is quantified based on his ability to discriminate between a control medium and a female-conditioned medium [12,13]. In particular, these studies have identified ascaroside pheromones as important in mate choice signaling due to their sex-specific production and effects on sexually-associated behaviors [14,15]. Female pheromones act as a male attractant in both hermaphroditic *C. elegans* [12] and gonochoristic *C. remanei* [16]. This female-based signaling appears to be related to the amount of sperm stored [16-18] and targets male-specific neurons [13,14,16,19,20]. Recent work has turned to male-produced ascarosides, showing that hermaphrodites exposed to male pheromones alone have a decrease in lifespan [21-24]. However, such male-produced pheromones do not appear to elicit a female mate choice response [12,16,22]. While ascarosides play a predominant role in mate choice, they also influence other population behaviors, thus necessitating a precise combination and concentration of small molecules for signaling specific to male-female interactions [11,14]. Since pheromone cues depend on such a precise mixture, accurate intersex communication assays necessitate a well-controlled environment where the concentration and diffusion of molecules is clearly defined and external signals are limited.

Microfluidic technology has proved to be an excellent method in which to study behavioral responses under precise environmental control [25-27]. Microfluidic devices scale on the nano-to micro-size and thus the small size of *Caenorhabditis* makes them suitable for manipulation within a microfluidic environment. Microfluidics offers many advantages over traditional plate-based assays, including better control of the concentration of molecules and their diffusion due to the laminar properties of microfluidics [28]. Several microfluidic devices have been designed to study behavioral responses such as chemotaxis, thermotaxis, and electrotaxis [25-27,29,30]. With respect to reproductive behavior, Chung et al. [29] found a behavioral response of individually isolated males when exposed to a uniform concentration of hermaphrodite-conditioned medium. In particular, they showed that males exposed to pre-conditioned medium spent more time performing sexually-associated behaviors than those exposed to a control medium. While their microfluidic device overcomes the issues of pheromone diffusion on agar plates, it cannot be used to study mate-choice searching patterns [as in 20,31] or direct choice comparisons between different attractants. Additionally, the pheromone signal contained in the pre-conditioned media is likely to decrease over the time required to study such locomotion patterns. Moreover, conditioned medium may be difficult to accurately reproduce as the concentration of small molecules will depend on the density of worms used to produce pheromones as well as the time spent in liquid culture. Given the limitations of exposure to a single pre-conditioned medium, the relationship between a pheromone signal and the elicited sexual response should be studied in real time. Given the limitations of exposure to a single pre-conditioned medium, the relationship between a pheromone signal and the elicited sexual response should be studied in real time.

We describe a new microfluidic paradigm, using the Pheromone Arena microfluidic device, that overcomes the current limitations of traditional plate-based assays and existing microfluidic technology to study sex-specific mate choice with constant exposure to pheromones produced in real time. We show that the Pheromone Arena allows for small-molecule communication alone without a decay of signal over time, thus validating its use for mate choice assays. Using the Pheromone Arena, we show that males are more attracted to virgin females than mated ones in both *C. remanei* and *C. elegans* with the degree of attraction being species-dependent. Additionally, we show that females rely on pheromone cues, though their preference is dependent on mating status.

## Materials and Methods

### Worm culture and strains

Wildtype *Caenorhabditis remanei* (strain EM464) and feminized *C. elegans* (strain JK574: *fog-2* mutation on the standard N2 laboratory background) were used in this study. Feminization of *C. elegans* is achieved by blocking self-sperm production in hermaphrodites, making them functionally female, and they will be referred to thusly. Use of a feminized *C. elegans* hermaphrodites allowed for direct comparisons between species as well as preventing any potential mate cue effects due to self-sperm [see 17,18]. Both strains were grown at 20°C on NGM-agar plates seeded with OP50 *Escherichia coli* bacteria following Brenner [32]. Synchronized cultures of stage 1 larvae were prepared by hypochlorite treatment of gravid females [33]. Larvae were matured to young adulthood in population densities of approximately 1,000 individuals. To maintain virgins, males and females were separated onto sex-specific plates of 40-50 individuals 40-45 hours post-larval stage 1. Mating plates of 25 females and 25 males were created at the same time. At the start of all the choice assays, day 1 adult virgins were 48 hours post-larval stage 1 and day 2 adults were 72 hours post-larval stage 1 (24 hours after separation to virgin or mating plates).

### Microfluidic device manufacturing

The Pheromone Arena (final design: v2.1; S1 File) was designed using CAD software (Vectorworks 2013 SP5, Nemetschek Vectorworks, Inc). Single layer devices were fabricated out of polydimethylsiloxane (PDMS) following soft lithography methods [34] and bonded to a glass microscopy slide following exposure to air plasma. Holes for connecting tubing were punched using a 1.25mm biopsy punch.

### Microfluidic set-up and experimental protocol

To avoid blocking flow, air bubbles were evacuated from each device using a vacuum chamber and replaced with M9 buffer. Worms were loaded into three chambers: ten worms each were loaded into the two upstream “signaling” chambers of the device and 20 to 25 worms were loaded into the single downstream “choic” echamber. To control for environmental and observational biases, worm combinations were alternatively loaded into the left and right upstream chambers.

Liquid was flowed through all inlets continuously, with the start of flow corresponding tothe beginning of an experiment. Flow was maintained at a constant rate using a pressurized air system (S1 Fig.). Specifically, air exiting a pressurized one gallon tank was regulated to 1.5 PSI. The air-line running from the tank was bifurcated to two tubing lines, each pressurizing a sealed 500mL bottle of M9. The bottle caps were modified to hold seven pieces of tubing: one for the air-line (terminating at the cap) and six liquid supply lines. The liquid tubing lines extended below the M9 surface, allowing liquid to flow out of the bottle and into the microfluidic device once air pressure was applied from the air-line. Tubing lengths were equal for each partition of the set-up to maintain equal flow through all lines. Pressure could be maintained for three hours off a single air tank.

The head position of each worm in the downstream chamber was counted every 30 minutes and recorded as being located under the left or right upstream chamber. Worms located in the filter separators were not counted. If worms were flushed out of the downstream chamber (and therefore the device), the assay continued without counting these worms and thus some replicates had a decrease in sample size over time. If worms climbed or were flushed into a different chamber than the one into which they were loaded, the experiment was terminated. Each choice combination was replicated multiple times (n ≥ 3) over multiple days (n ≥ 2). Day 2 adult worms were used for the male choice assays. Day 1 adult worms were used for female choice assays, except when comparing virgin versus mated female choice, which required using day 2 adult females. All data have been made available (S2 File).

### Statistical analyses

Data were analyzed in R v3.2.1 (R Core Development Team 2015). Replicates were pooled for each choice assay by time point. An equality of proportions test was performed for each assay individually to determine if: i) males were more attracted to virgin females over mated females or ii) females were more attracted to virgin males over virgin females. The null hypothesis was that of no choice (using a probability of success = 0.5). A chi-square test for homogeneity was preformed to determine if the proportion of males or females choosing virgin females or males, respectively, was equal across species combinations.

## Results

### Microfluidic design

The Pheromone Arena has three sequential components: three inlets with each with a distribution loading network, a main arena, and a single high resistance outlet (Fig. 1A). The arena area is further divided into three physically distinct chambers. The chambers all have a pillar-array to facilitate the natural, sinusoidal movement of worms [35,36]. The two upstream chambers hold the pheromone signalers. Two versions of the loading distribution networks were created to accommodate the size differences between males and females at day 2 of adulthood. Specifically, female signalers have a wider distribution network, while male signalers have a narrower distribution network coupled with the removal of the first two rows of pillars to prevent males from climbing out of the chamber (Fig. 1B). The upstream chambers are separated from the single, large downstream chamber by a very fine filter (Fig. 1C). This filter allows for small molecules to pass, but rarely can worms pass through. Worms making a choice were loaded into the downstream chamber by a third inlet. Due to laminar flow, two pheromone environments are created in the downstream chamber that mirror the signaling pair in the upstream chambers (Fig. 1D). All the chambers were greater than one-by-one worm length, allowing for free movement of individuals without density effects (Fig 1E).

**Fig 1.**
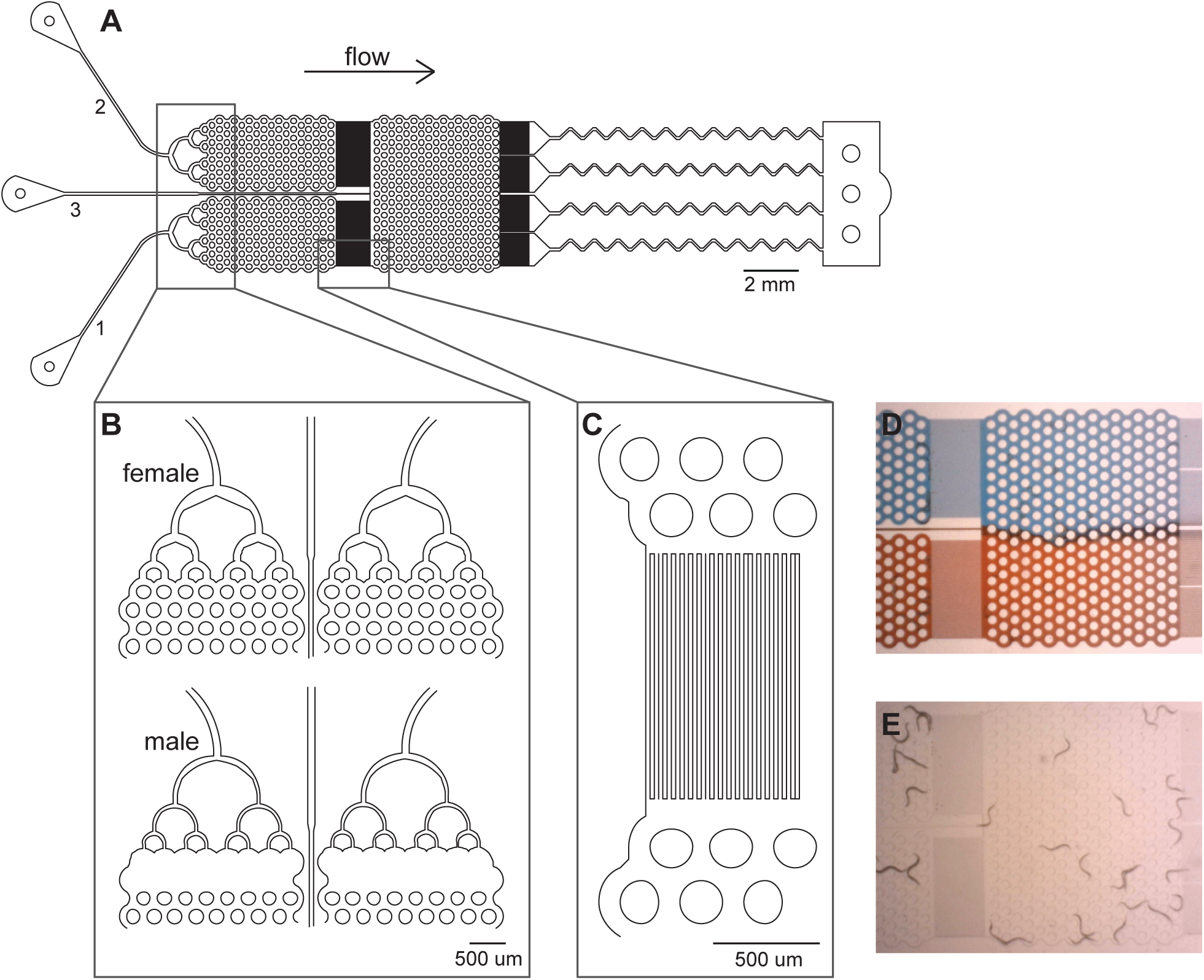
Microfluidic Pheromone Arena for mate choice assays. (A) Blueprint for the Pheromone Arena (v2.1; S1 File). The three inlets correspond to the three worm chambers: inlets 1 and 2 connect to the upstream chambers and inlet 3 connects to the downstream chamber. All the chambers have a pillar-array (shown as circles) spaced 100um apart to allow for natural worm movement. An 18um filter separates the downstream chamber from the upper two chambers and from the outlet. Extra resistance was added to the outlet distribution to decrease the overall flow rate. (B) Close-up of the loading distribution networks for the upstream chambers. The distribution network used depended on whether males or females were the pheromone signalers. (C) Close-up of the filter separating chambers. (D) Visualization of how the laminar flow dynamics create two distinct environments in the downstream chamber using red and blue dyes. (E) Visualization of worms in the device. After being loaded, worms move freely throughout their chamber without passing into another chamber.

No inherent biases were measured in the movement of worms in the downstream chamber. In particular, when no pheromone cue was present, males were equally likely to move to the left of right sides of the chamber in a random fashion (p = 0.45, n = 24 replicates).

## Males are attracted to virgin females

Virgin *C. remanei* and *C. elegans* males were assayed for their ability to discriminate between conspecific virgin and mated females. Male choice was measured approximately every 30 minutes for 3 hours. Across all replicates, male choice varied little after 60 minutes (S2 Fig.). Therefore, we used data from this time point as a single comparative measure of male choice across assays.

Males were more attracted to virgin females than mated females in both *C. remanei* (proportions test: *χ*^2^ = 46.2, d.f. = 1, p < 0.0001, 95% C.I. of virgin attraction = 72.6-87.1%) and *C. elegans* (proportions test: *χ*^2^ = 4.15, d.f. = 1, p = 0.04, 95% C.I. of virgin attraction = 50.3-64.8%), though the ability to discriminate was much weaker in *C. elegans* (Fig. 2). To better understand this marked difference in male sensitivity, male choice was assayed for the ability to discriminate between virgin and mated heterospecific females. In both heterospecific crosses males were more attracted to virgin females than mated females (*C. remanei* male proportions test: *χ*^2^ = 42.0, d.f. = 1, p < 0.0001, 95% C.I. of virgin attraction = 66.1-78.7%; *C. elegans* male proportions test: *χ*^2^ = 24.2, d.f. = 1, p < 0.0001, 95% C.I. of virgin attraction = 64.1-80.9%). For example, *C. remanei* males chose virgin *C. elegans* females more successfully than in the conspecific *C. elegans* assay. Similarly, *C. elegans* males had a much high ability to discriminate virgins when they were presented with *C. remanei* females. The strength of attraction to virgins for both heterospecific assays was between that of the conspecific assays, suggesting species-dependent pheromone effects in both males and females (*χ*^2^ = 22.0, d.f. = 3, p < 0.0001).

**Fig 2.**
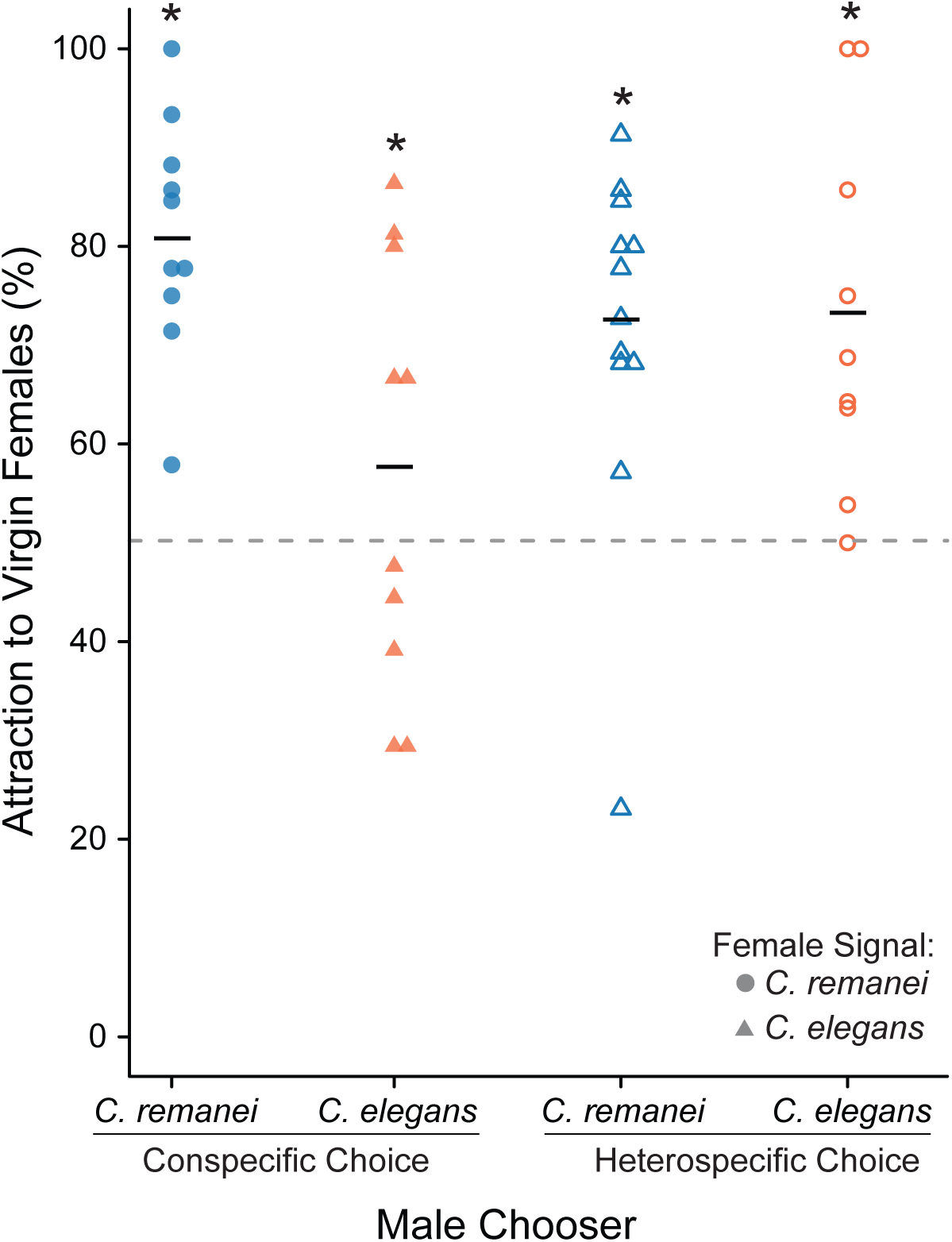
Males are more attracted to virgin females than mated females within and between species. Virgin, day 2 adult *C. remanei* males (blue) and *C. elegans* males (orange) were given a choice between virgin and mated female pheromone (*C. remanei* females shown as circles and *C. elegans* females shown as triangles). Each replicate is represented as an individual point and the mean attraction to virgin females is given by the horizontal bar. Conspecific assays are shown as solid point and heterospecific assays as open points. The null hypothesis of no choice is given by the dashed line. In each assay males were more attracted to virgin females than mated ones. However, the strength of male attraction was dependent on the species of both the chooser and the signaler (Test of homogeneity across assays: *χ*^2^ = 22.0, d.f. = 3, p < 0.0001). Asterisks denote a significant proportions test.

## Females choose male pheromone over those of from females

Virgin *C. remanei* and *C. elegans* females were assayed for their ability to discriminate between conspecific virgin males and females. Female choice increased slightly over time, but was consistent by 60 minutes, again making this time point a reflective measure of overall female choice (S3 Fig.). Interestingly, *C. remanei* and *C. elegans* followed a similar trend in choice of males over time, though at very different magnitudes. Specifically, female *C. remanei* chose conspecific male pheromones over female pheromones (proportions test: *χ*^2^ = 14.8, d.f. = 1, p < 0.001, 95% C.I. of male attraction = 58.0-74.0%) (Fig. 3). When females were given a choice between male pheromone and no pheromone, they still chose the male pheromone more than expected by chance (*χ*^2^ = 7.19, d.f. = 1, p < 0.01), suggesting that females are in fact attracted to males and not simply repulsed by other females. However, *C. elegans* females showed no clear differentiation between conspecific males and females (proportions test: *χ*^2^ = 0.563, d.f. = 1, p = 0.45, 95% C.I. of male attraction = 45.0-61.8%). Moreover, the heterospecific assays also showed a lack of discrimination between male and female pheromones, suggesting that female choice may be species-specific, or at least very weak at best within *C. elegans*.

**Fig 3.**
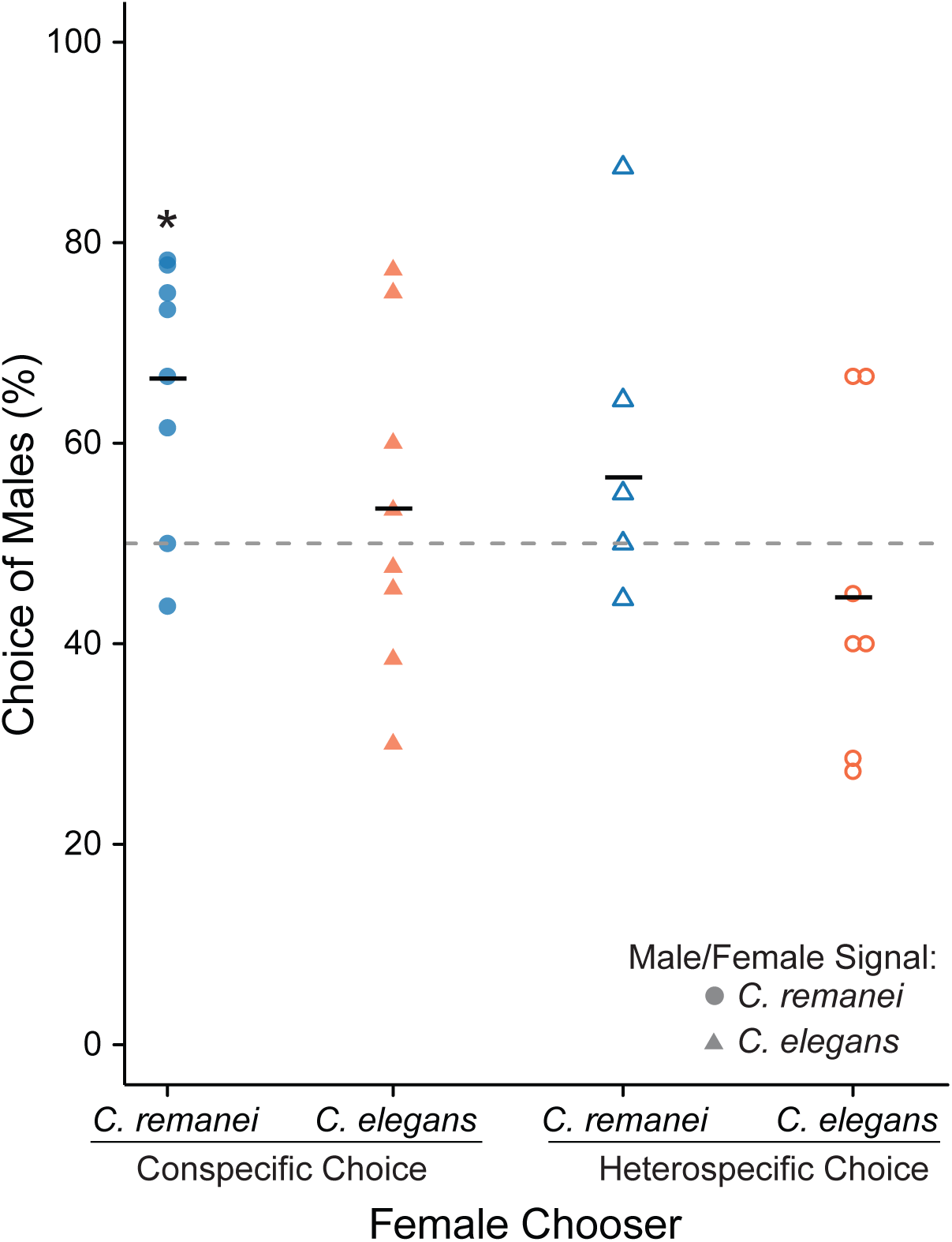
Female choice of virgin males over virgin females is species-specific. Virgin, day 1 adult *C. remanei* females (blue) and *C. elegans* females (orange) were given a choice between virgin male and female pheromone (*C. remanei* females shown as circles and *C. elegans* females shown as triangles). Each replicate is represented as an individual point and the mean attraction to males is given by the horizontal bar.Conspecific assays are shown as solid point and heterospecific assays as open points. The null hypothesis of no choice is given by the dashed line. Only when *C. remanei* females were given a choice of conspecifics were they more attracted to male pheromone than female pheromone (*χ*^2^ = 14.8, d.f. = 1, p < 0.001). All other comparisons failed to reject the null hypothesis of no female choice. Asterisks denote a significant proportions test.

**Fig 4.**
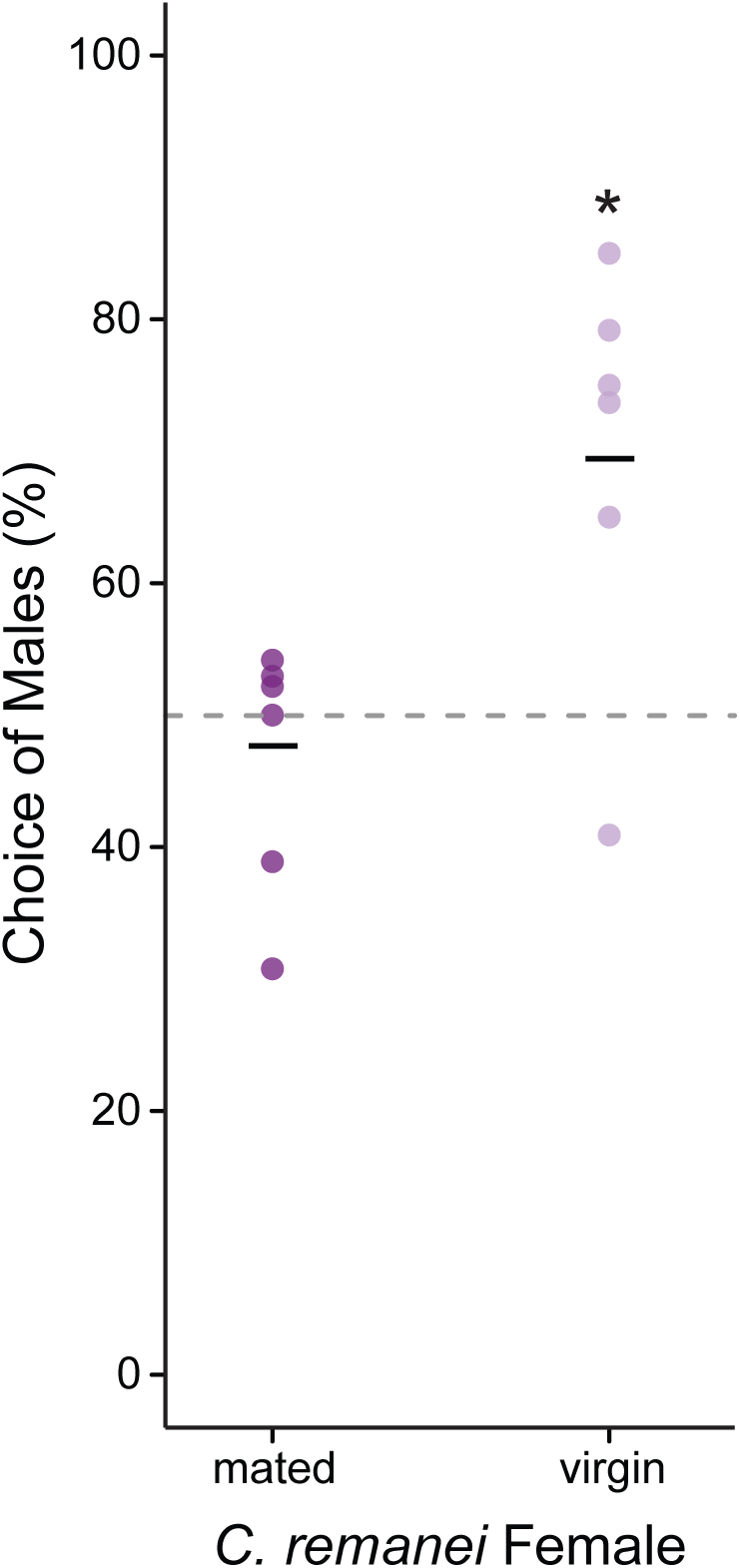
Female attraction depends on mating status. Virgin (light purple) and mated (dark purple) day 2 adult *C. remanei* females were given a choice between virgin day 1 adult *C. remanei* male and female pheromones. Each replicate is represented as an individual point and the mean attraction to males is given by the horizontal bar. The null hypothesis of no choice is given by the dashed line. Virgin females showed a strong preference for males over females (*χ*^2^ = 17.5, d.f. = 1, p < 0.0001), while mated females displayed no choice (p = 0.70). Asterisks denote a significant proportions test.

Additionally, we examined if female choice was dependent a female mating status. The female choice assay was replicated using mated and virgin *C. remanei* females. Virgin females showed a strong preference for virgin male over female pheromones (proportions test: *χ*^2^ = 17.5, d.f. = 1, p < 0.0001, 95% C.I. of male attraction = 60.3-69.4%). However, mated females showed no obvious preference (proportions test: *χ*^2^ = 0.150, d.f. = 1, p = 0.70, 95% C.I. of male attraction = 38.0-57.5%).

## Discussion

Mate choice is ubiquitous across metazoans with sexual reproduction. Understanding the signal-receiver dynamics of mate choice beyond phenotypic traits – such as pheromone signals – provides valuable information on how individuals discern high quality mates with a propensity to mate. Here we proposed a new microfluidic paradigm to quantify pheromone communication in nematodes. Our design allows for real time isolation of small molecules in a controlled environment as well as allowing for the natural searching and choice behaviors of receivers over time. While this design is not the first to use pillared-arenas [35-37], to the best of our knowledge no other worm-specific microfluidic devices have such a physically separated, sequential arena design. Moreover, the Pheromone Arena expands on previous worm choice devices [29,30] by allowing for natural searching behaviors in addition to measuring overall choice. Specifically, we examined male and female mate choice in *C. remanei* and *C. elegans* using a combination of conspecific and heterospecific assays to determine how mating status affects attraction. This study is the first use this combinatorial design coupled with real time pheromone signaling and spatial complexity.

The conspecific male choice results support previous studies [16,18] in showing that virgin males are strongly attracted to virgin females over mated females in both *C. elegans* and *C. remanei*. Interestingly, males from these species did not discriminate between female mating types with the same intensity [16]. In particular, *C. elegans* males had a reduced ability to discern virgin females from mated ones. However, when *C. elegans* males were presented with pheromones from *C. remanei* females, male attraction to virgins increased. Similary, *C. remanei* males could distinguish between virgin and mated *C. elegans* females better than *C. elegans* males. Therefore, there appears to be a decrease in both female signal intensity as well as male receptor capability in *C. elegans*, leading to an overall decrease in mate choice ability. This diminution of choice is likely a result of the independent lineage transition to self-fertilizing hermaphrodism in *C. elegans*. Since fertilization is predominantly by selfing and males are rare within populations, sexual selection – apparently including mate recognition dynamics – is greatly reduced.

This hypothesis could be further tested by altering the number of signalers in the upstream chambers. The pheromone arena allows for exact control over the number of worms producing pheromone and, given the constant flow dynamics, altering the number of worms would in effect alter the concentration of pheromone signal in the downstream chamber. Previous studies have used various concentrations of hermaphrodite-conditioned media and found different results in male attraction [16,18]. Our assays used the same number of females in both upstream chambers, however, modulating the number of females in each upstream chamber is promising future work.

Previous work has shown that females are attracted to isolated, male-produced ascaroside cues [15,38], however, this result has not been replicated when the signal is produced *in vivo*. We took a novel approach comparing female discrimination between male and female produced pheromones to determine if females are truly attracted to males or are simply repulsed by the presence of a high number of other females. A discernable choice of males over females was measured in the *C. remanei* conspecific assays. Additionally, females were attracted to male pheromone over a no pheromone control, suggesting this choice measured is true attraction to males. Despite making a choice, the intensity with which females choose males was much weaker than seen for the male choice assays. This weak attraction could potentially explain why previous plate-based assays – where male signals can be lost by diffusion or mixed with other signals from the environment – could not measure any female choice of males [13,16]. However, *C. elegans* females made no choice in the conspecific assays as was also seen for both heterospecific assays, suggesting species-specific effects. Together these results suggest that when sexual selection is strong, as in *C. remanei*, males take a more active searching approach, such that females are predominantly signalers, while males are active receivers and searchers. This signal-receiver dynamic is in somewhat of a sex-role reversal from traditional sexual selection models of male signaling and female receiving, though still consistent with anisogamy.

We further examine female choice based on a female’s mating status. Mating status is known to influence remating behavior in many species, such that mated females – or females with sperm – typically have a lower propensity to remate [39,40]. Moreover, mated females will run away from males to avoid remating [41]. These observation is consistent with our results as virgin females strongly preferred male pheromones while mated females made no choice between male and female pheromones.

While the Pheromone Arena is both an innovative and effective tool, there are several limitations to its use. Namely, the difference in size between older adult males and females poses a problem for intersex comparisons. Additionally, the design could be improved by limiting worms from reaching other chambers through the filter separators. This cross-chamber movement was particularly an issue with males as they have a small diameter and a highly-developed searching behavior that leads them to attempt to crawl through the filters to reach the virgin females. However, the filter between the upstream and downstream chambers is currently at the lower limit of what can accurately be manufactured using PDMS-based microfluidics and thus a significant design change would be required to prevent this tenacious behavior.

Despite these limitations, the Pheromone Arena was able to reproduce previously seen sexual behavior responses and go further into the study of species-specific sexual attraction in both males and females. In the future, this type of device could be used to study the effect of density or sex ratio on sexual attraction. Moreover, it would be possible to modify these devices to allow food delivery and perform longer term assays or study the influence of food availability on sexual attraction. Such assays would benefit from being coupled with an automated system [see 42] to obtain a more accurate counting via recordings and to facilitate high-throughput experiments.

## Acknowledgements

We would like to thank Nadine Timmermeyer for prototyping Pheromone Arena v1.0 and Ben Blue for assistance with microfluidic manufacturing and advice on the pressurized air system.

## Supporting Information

**S1 File. Pheromone Arena microfluidic design.** The Pheromone Arena has two versions to account for the difference in size between day 2 adult males and females: male choice of female signalers and female choice of male signalers. In the male choice version, the upstream loading distribution network is sized at 60um to account for the larger diameter of females, while the downstream chamber distribution channel has a final constriction size of 35um to prevent males from climbing back out of the downstream chamber. The female choice version has a smaller upstream distribution network (40um) and a larger downstream chamber distribution (60um). Additionally, in the female choice version the two upstream chambers have the first two rows of pillars removed, again to prevent males from climbing out of the device. Both versions have increased resistance added to the outflow to decreased the overall flow rate through the device. These blueprints are accessible using CAD software. A master height of 65um is recommended.

**S2 File. Data.** Raw data for all male choice, female choice, and control experiments.

**S1 Fig. Pressurized air flow set-up.** A pressurized air system was used to maintain a constant flow rate through the microfluidic devices. A one gallon air tank was regulated to 1.5 PSI was sufficient to run experiments for up to 3 hours. The air-line running from the tank was bifurcated to pressurize two sealed 500mL bottles of M9 buffer. The bottle caps were modified to supply six liquid lines, which connected to two Pheromone Arenas, as well as hold the air-line, which terminated at the cap. The tubing lengths were equal for each partition of the set-up to maintain equal flow through all lines. The Pheromone Arena was kept on a confocal microscope at 20°C for the duration of each experiment. This figure was modified with permission from Stephen Banse.

**S2 Fig. Virgin male choice over time.** Virgin, day 2 adult *C. remanei* males were given a choice between virgin and mated conspecific female pheromone (blue) and *C. elegans* males were given a choice between virgin and mated conspecific female pheromone (orange). The weighted means and standard error are plotted over time. By 60 minutes into the experiment males had made a consistent choice (down triangle). The null hypothesis of no choice is given by the dashed line. In each assay males were more attracted to virgin females than mated ones.

**S3 Fig. Virgin female choice over time.** Virgin, day 1 adult *C. remanei* females were given a choice between virgin conspecific male and female pheromones (blue) and *C. elegans* males were given a choice between virgin conspecific male and female pheromones (orange). The weighted means and standard error are plotted over time. The null hypothesis of no choice is given by the dashed line. Only *C. remanei* females made a measurable choice of male pheromone over female pheromone.

